# Transcriptional modulation during photomorphogenesis in rice seedlings

**DOI:** 10.1101/2023.09.08.556755

**Authors:** Parul Gupta, Pankaj Jaiswal

**Author notes:** **Correspondence:** Prof. Pankaj Jaiswal, Department of Botany and Plant Pathology, Oregon State University, Corvallis, Oregon, USA.

## Abstract

Light is one of the most important factors regulating plant gene expression patterns, metabolism, physiology, growth, and development. To explore how light may induce or alter transcript splicing, we conducted RNA-Seq-based transcriptome analyses by comparing the samples harvested as etiolated seedlings grown under continuous dark conditions vs. the light-treated green seedlings. We identified 14,766 differentially expressed genes, of which 4369 genes showed alternative splicing. We observed that genes mapped to the plastid-localized methyl-erythritol-phosphate (MEP) pathway were light-upregulated compared to the cytosolic mevalonate (MVA) pathway genes. Many of these genes also undergo splicing. These pathways provide crucial metabolite precursors for the biosynthesis of secondary metabolic compounds needed for chloroplast biogenesis, the establishment of successful photosynthetic apparatus, and photomorphogenesis. In the chromosome-wide survey of the light-induced transcriptome, we observed intron retention as the most predominant splicing event. In addition, we identified 1709 novel lncRNA transcripts in our transcriptome data.

## INTRODUCTION

Light is an essential growth factor for sustaining autotrophic plant life. The quality, quantity, direction, and length of light exposure affect plant growth and development ^1,2,3^. The natural light-dark cycle maintains the carbon and nitrogen metabolism of rice plants ^4^. Light induces transcriptional reprogramming in various plant species ^4–12^ via an array of photoreceptors ^13,14^. The photoreceptors phytochrome (red light) and cryptochrome (blue light) are involved in light signaling and photomorphogenesis ^15,16^. Genes encoding phytochrome interacting factors (PIFs) promote skotomorphogenesis and development in the dark ^17–19^. Light-mediated gene expression modulation is also triggered by translational enhancement of the preexisting mRNA pool instead of an enhanced transcription rate ^20,21^.

Intron splicing in multiexonic mRNA is a post-transcriptional regulatory process that often produces different mRNA isoforms transcribed from a single gene locus, thus contributing to proteome plasticity. Light-induced alternative splicing (AS) was observed in ~10% of the protein-coding genes of *Arabidopsis* and *Physcomitrella patens* ^22,23^. A flash of light applied in the middle of the dark or nighttime is sufficient to induce splicing ^24^. In contrast to animals, where the most common AS event is exon skipping (ES), intron retention (IR) is the most common AS event in rice ^25^, Arabidopsis ^25,26^, and Poplar ^27^. It is now well established that transcriptome modulation via AS is vital for plant growth, development, and stress response ^28–30^.

Compared to the protein-coding genes, long non-coding RNAs (lncRNAs) are transcripts >200 bp in length that do not have protein-coding potential. lncRNAs are classified into categories: (i) sense lncRNAs, (ii) antisense lncRNAs, (iii) intergenic RNAs (lincRNAs), (iv) intronic RNAs, and (v) bidirectional lncRNAs ^31^. In addition to the numerous studies coupling gene expression and AS studies on protein-coding genes, it is now known that lncRNAs also play a role in regulating gene expression through transcriptional, post-transcriptional, and chromatin remodeling mechanisms ^31–34^. In rice, lncRNAs regulate biological processes, such as ovule development and female gametophyte abortion ^35^, sexual reproduction ^36^, and stress response ^37,38^. However, the role of light in regulating lncRNAs is not well studied. *Arabidopsis* non-coding RNA *HID1* is a known positive regulator of photomorphogenesis in continuous red light ^39^.

Rice is a global staple crop and is a model for studying crop genomics. The transition from skotomorphogenesis under dark conditions to photomorphogenesis under light exposure is critical for seedling survival and requires precise control of gene expression by different regulatory mechanisms. Results from our rice study show that exposure to light alters the expression and splicing of a wide array of protein-coding genes but not so much for the non-coding lncRNA genes.

## MATERIALS & METHODS

### Plant material, growth conditions, and treatment

Seeds of rice (*Oryza sativa spp. japonica* cv. Nipponbare) were grown and processed for the experiment by following the growth conditions and sampling described previously ^40^. Seeds were sown in the dark and germinated on day 2. After sowing, these germinated seedlings grew in the dark for 8 days (8DD). At the end of day 8, three biological replicates of the dark-grown etiolated shoots were harvested. The remaining dark-grown seedlings were exposed to continuous white light at 120 µmol/m2/sec (measured at the soil surface) for 48 hours or 2 days (days 9 and 10 after sowing). The shoots of three biological replicates of light-treated green-colored seedling samples (8DD-2LL) were harvested at the end of day 10. Harvested samples were frozen using liquid nitrogen and stored at −80°C until further processing. Throughout this report, 8DD-treated plants are called dark samples, and 8DD-2LL light-treated plants are called light samples. Data analysis is described for light regulation.

### Sample preparation and sequencing

Total RNA from frozen tissue samples was extracted using RNA Plant reagent (Invitrogen Inc., USA) and RNeasy kits (Qiagen Inc., USA) and treated with RNase-free DNase (Life Technologies Inc., USA) according to the manufacturer’s protocol. The total RNA quality and concentration were determined using an ND-1000 spectrophotometer (Thermo Fisher Scientific Inc., USA) and Bioanalyzer 2100 (Agilent Technologies Inc., USA). PolyA-enriched mRNA libraries were prepared from three biological replicates of dark and light samples using the TruSeqTM RNA Sample Preparation Kits (v2) and sequenced as 51-bp single-end reads using the Illumina HiSeq 2000 instrument (Illumina Inc., USA) according to the manufacturer’s protocol at the Center for Genomic Research and Biocomputing, Oregon State University. The strand-specific sequencing reads and metadata were deposited at EMBL-EBI ArrayExpress (accession number E-MTAB-5689).

### RNA-seq data analysis

The generation of FASTQ files from the RNA-Seq sequences was performed by CASAVA software v1.8.2 (Illumina Inc.). Sequence reads were filtered and trimmed for low quality at a score of 20 using Sickle v1.33 ^41^. Clean, high-quality reads from each sample and replicates were aligned to the *Oryza sativa japonica* cv Nipponbare reference genome (IRGSP-1.0.31) using TopHat v2.1.1 ^42^. Mapped reads were assembled using Cufflinks, and the reference-guided assembled transcripts from each replicate were merged using Cuffmerge ^43^. Assembled transcripts were compared to the reference genome annotation using Cuffcompare. The RSEM software package estimated normalized baseline expression from the aligned sequence reads ^44^. For differential gene expression analysis, read count data obtained from RSEM were used in EBSeq ^45^. Differentially expressed (DE) genes were filtered based on the false discovery rate corrected P value ≤0.05.

### Functional annotation

We carried out the Gene Ontology (GO) enrichment analysis tool provided by the GO consortium ^46^ to determine the biological roles played by the enriched gene set. Plant pathway enrichment analysis was done by mapping the DE genes using the Plant Reactome analysis tool ^47,48,49^ (http://plantreactome.gramene.org/PathwayBrowser/#TOOL=AT).

### Alternative splicing analysis

Splicing events in the transcripts from the samples were identified by the SpliceGrapher v0.2.5 pipeline ^50^. Sequence reads from each sample, and replicates were aligned to the reference rice genome. Splice site-specific classifiers were built using build_classifiers.py script using canonical (GT) and noncanonical (GC) donor sites and acceptor site (AG) for *Oryza sativa* genome annotation version 31 (Oryza_sativa.IRGSP-1.0.31). Read alignments in SAM format from each replicate were used as input for SpliceGrapher’s sam_filter.py script to filter out false-positive sites. SpliceGrapher Python scripts were used for the generation of depth files (sam_to_depths.py), splice graph prediction (predict_graphs.py), generating statistics (splicegraph_statistics.py) from a set of splice graphs, gene-by-gene summary (genewise_statistics.py) of splicing events and splice graph visualization (plotter.py). The Realignment pipeline was used to construct putative transcripts from unresolved exons with sufficient coverage from the alignments ^50^. In the following steps, IsoLasso ^51^, an extension of the SpliceGrapher workflow, was used to predict novel splicing events.

### Prediction of long non-coding RNA (lncRNA)

All transcripts annotated as intergenic transcripts, intron transcripts, antisense exon transcripts overlapping the reference exons, and antisense intron transcripts overlapping the reference introns were considered potential lncRNA candidates. Transcript sequences of length <=200 nucleotides were filtered out, and the gffread function of Cufflinks was used to extract fasta sequences of potential lncRNA transcripts from the gtf file. CPC2 ^52^ was used to predict the coding potential of transcripts. Predicted lncRNAs were scanned by InterProScan ^53^ to ensure the absence of protein-coding domains. To identify novel lncRNAs, a BLASTn ^54^ search was performed against a custom BLAST database generated using rice lncRNAs downloaded from the CANTATAdb2.0 (http://cantata.amu.edu.pl), PNRD ^55^, GreeNC ^56^, and RiceLNCPedia ^57^ databases. Secondary structures for the lncRNA molecules were predicted by the RNAfold software ^58,59^.

## RESULTS

### Light-mediated differential gene expression during photomorphogenesis

To explore the transcriptome modulation in rice in response to light, we sequenced the strand-specific poly-A enriched RNA fraction isolated from three biological replicates of rice plants grown under dark and light exposure conditions (see methods). A total of ~38 million and ~42 million high-quality reads were generated from the dark and light-treated samples, respectively. More than 92% of reads from each sample aligned to the rice reference genome (Supplementary Table S1). We found 38,642 genes showed baseline normalized expression in the samples, of which 33,943 genes expressed in the dark vs 35,772 genes that expressed under light, respectively. Differential expression analysis identified 14,766 light-regulated genes (Figure 1, Supplementary Figure S1, Supplementary Table S2).

**Figure 1:**
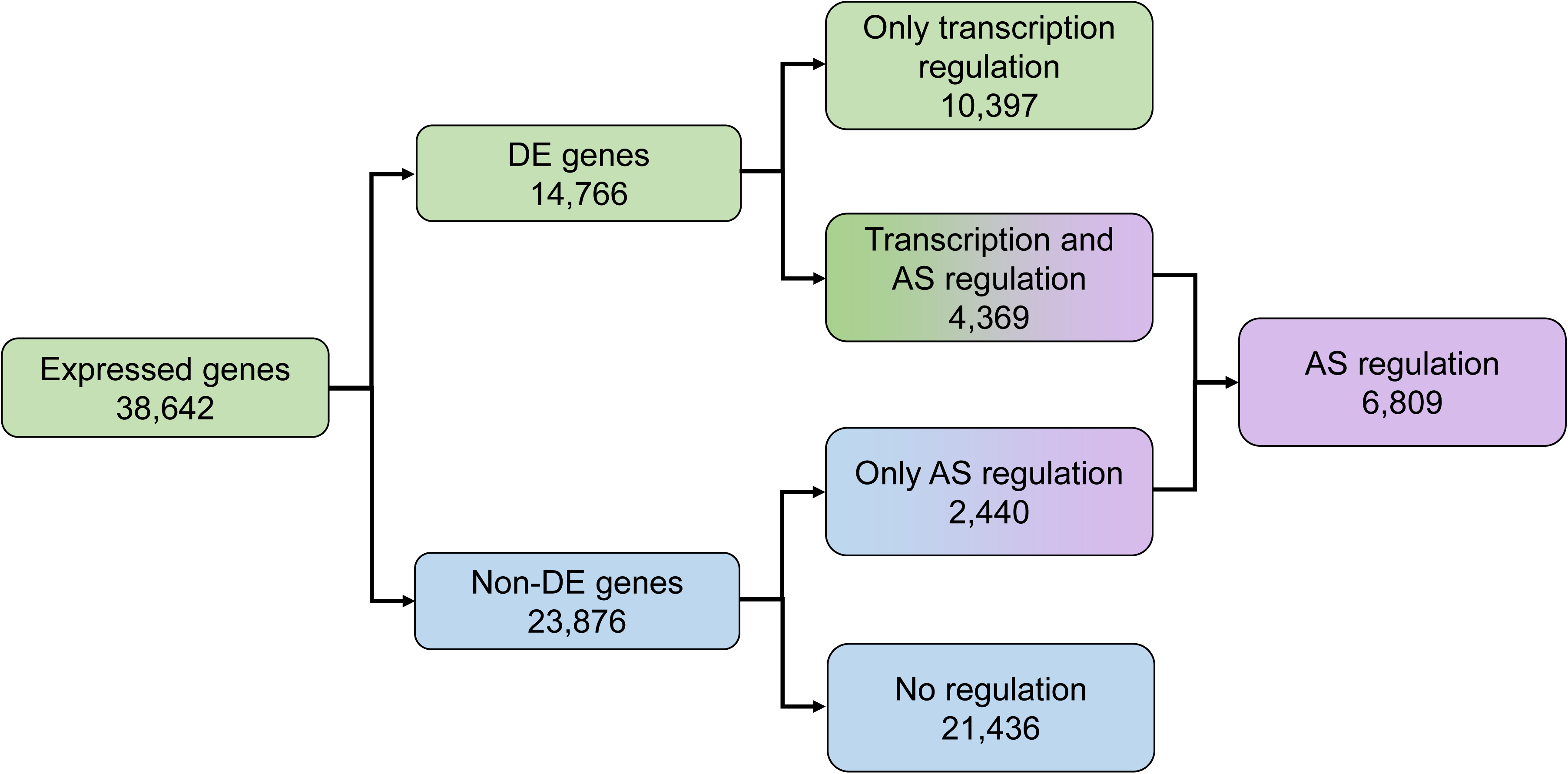
Summary of differential gene expression and transcript splicing observed in rice seedling shoots undergoing photomorphogenesis.

### Gene function and pathway enrichment analyses

Gene Ontology-based functional annotation analysis of the light-regulated differentially expressed gene set (Supplementary Figure S2) revealed enrichment for biological processes (BP), such as chlorophyll and cell wall biosynthesis, besides photosystem light reaction pathways and ribosome assembly. Dominant molecular functions (MF) were rRNA binding, cytoskeletal motor activity, oxidoreductase and metallopeptidase activities, translation elongation, and chaperone binding. As expected, chloroplast, cytoskeleton, ribosomes, peroxisomes, and nucleus were the major cellular component sites of activity.

Pathway enrichment analysis using the Plant Reactome pathway analysis tool ^60^ mapped 475 light-upregulated genes to 243 pathways (Supplementary Table S3). A total of 164 pathways overlapped between light-up and light-downregulated genes. Most pathways showed higher mappings to light-upregulated genes, except hormone auxin and brassinosteroid signaling and reproductive structure development (seed). Pathways unique to light-upregulated genes include those for the biosynthesis of photosynthesis components chlorophyll, carotenoid, and phylloquinone, hormones like gibberellin, auxin, abscisic acid, *etc*. The pathways with unique mapping to light-downregulated genes include polar auxin transport, mevalonate (MVA) pathway, circadian clock, salicylic acid metabolism and signaling, reproductive plant part development, root-specific gene network of NAC10 transcription factor (Supplementary Figure S3). To our surprise, we observed enrichment of the

### Identifying light-regulated transcription factors

To identify the light-regulated transcription factors (TF), a list of rice TFs was downloaded from the Plant Transcription Factor Database ^61^ and searched against the DE genes. We found 429 light-upregulated and 498 light-downregulated TFs (Supplementary Table S4). We found WRKY, NAC, and orphans were the most abundantly expressed TFs in light, compared to many light-downregulated bHLH, bZIP, and C3H gene family members (Table 1). Highly upregulated (fold change ≥10) TFs belong to the MYB, AP2-EREBP, WRKY, Orphans, NAC, MADS, and bHLH gene families; however, down upregulated TFs belong to the AP2-EREBP, C2C2-CO-like, C3H, HB, and NAC gene families (Supplementary Table S4). To investigate whether TFs targeted the MVA and MEP pathway genes, we surveyed the list of TFs and their potential targets identified by the Plant Transcription Factor Database. We identified 23 TFs that potentially bind to the promoter region of 6 MVA pathway genes. Two MVA pathway genes, hydroxymethylglutaryl-CoA synthase (*HMGS*; Os08g0544900) and 3-hydroxy-3-methylglutaryl coenzyme A reductase (*HMGR*; Os08g0512700), were targeted by bZIP and AP2-EREBP factors, respectively, whereas the mevalonate 5-diphosphate decarboxylase (*MDD*; Os02g0109100) gene was a target of by 13 AP2-EREBPs, one C3H protein, and one C2H2 protein (Supplementary Table S5). None of the TFs we found bind to the promoter of MEP pathway genes.

**Table 1:** Light-regulated transcription factor gene families and their members.

| Gene family | Upregulated | Downregulated | Gene family | Upregulated | Downregulated |
| --- | --- | --- | --- | --- | --- |
| ABI3VP1 | 6 | 4 | GRF | 1 | 2 |
| Alfin-like | 1 | 4 | HB | 21 | 24 |
| AP2-EREBP | 23 | 32 | HSF | 3 | 10 |
| ARF | 8 | 8 | LIM | 2 | 1 |
| ARR-B | 1 | 4 | LOB | 2 | 1 |
| BBR-BPC | 0 | 1 | MADS | 6 | 5 |
| BES1 | 0 | 3 | mTERF | 9 | 4 |
| bHLH | 19 | 27 | MYB | 28 | 14 |
| BSD | 1 | 6 | MYB-related | 16 | 25 |
| bZIP | 10 | 27 | NAC | 21 | 19 |
| C2C2-CO-like | 4 | 7 | OFP | 2 | 1 |
| C2C2-Dof | 5 | 4 | Orphans | 20 | 10 |
| C2C2-GATA | 7 | 4 | PBF-2-like | 1 | 0 |
| C2C2-YABBY | 3 | 1 | PLATZ | 1 | 4 |
| C2H2 | 14 | 19 | RWP-RK | 4 | 2 |
| C3H | 16 | 24 | S1Fa-like | 1 | 1 |
| CAMTA | 0 | 3 | SBP | 3 | 5 |
| CCAAT | 9 | 11 | Sigma 70-like | 4 | 0 |
| CPP | 1 | 4 | SRS | 1 | 0 |
| CSD | 0 | 1 | TAZ | 1 | 1 |
| DBP | 2 | 3 | TCP | 4 | 6 |
| E2F/DP | 2 | 1 | Tify | 6 | 2 |
| EIL | 1 | 1 | Trihelix | 3 | 9 |
| FAR1 | 5 | 10 | TUB | 2 | 5 |
| FHA | 7 | 6 | ULT | 1 | 0 |
| G2-like | 9 | 5 | VOZ | 0 | 2 |
| GeBP | 0 | 3 | WRKY | 22 | 10 |
| GRAS | 8 | 7 | zf-HD | 5 | 3 |
| **Other transcriptional regulators** |  |  |  |  |  |
| Gene family | Upregulated | Downregulated | Gene family | Upregulated | Downregulated |
| ARID | 2 | 1 | PHD | 12 | 22 |
| AUX/IAA | 3 | 19 | Pseudo ARR-B | 2 | 1 |
| Coactivator p15 | 0 | 1 | RB | 0 | 1 |
| DDT | 3 | 0 | Rcd1-like | 0 | 2 |
| GNAT | 12 | 8 | SET | 11 | 6 |
| HMG | 4 | 5 | SNF2 | 11 | 9 |
| Jumonji | 4 | 5 | SOH1 | 0 | 2 |
| LUG | 0 | 2 | SWI/SNF-BAF60b | 4 | 4 |
| MBF1 | 0 | 1 | SWI/SNF-SWI3 | 2 | 1 |
| MED6 | 0 | 1 | TRAF | 7 | 11 |

### Transcript splicing during photomorphogenesis

To study how light exposure modulates transcript splicing (AS) events in the rice seedling shoots, we analyzed the transcriptome data by using SpliceGrapher ^50^. We created highly accurate splice site classifiers for rice with canonical (GT-AG) and semi-canonical (GC-AG) splice sites to filter the splice junctions in the sequence reads aligned to the reference genome. After removing the false positive splice sites, we predicted chromosome-wise splice graphs from the two transcriptome sample datasets. In the dark-treated dataset, 63.9% were true positive junctions, with 5.5% novel splice sites. In the light-treated dataset, 64.3% of splice junctions were true positives, of which 5.8% were predicted as novel sites. We observed 6214 spliced genes with 9685 splicing events in the dark samples compared to the 6809 spliced genes with 10432 splicing events in the light samples (Figure 1, Table 2). The highest number of spliced genes and events were on chromosome 1, and the lowest was on chromosome 10 (Table 2). Intron retention (IR) was the most prevalent type of splicing event in both samples, followed by exon skipping (ES). Alternative 3’ splicing (Alt.3’) was the least common event. To resolve ambiguous combinations of donor and acceptor splice sites, we realigned the sequenced reads from the two samples to the putative transcripts and resolved the novel exons to generate splice graphs. Light-induced splicing in 2162 unique genes compared to 1567 unique genes that undergo splicing in the dark. Of the 4647 spliced genes shared between the two samples, 165 genes displayed differential splicing events.

**Table 2:**
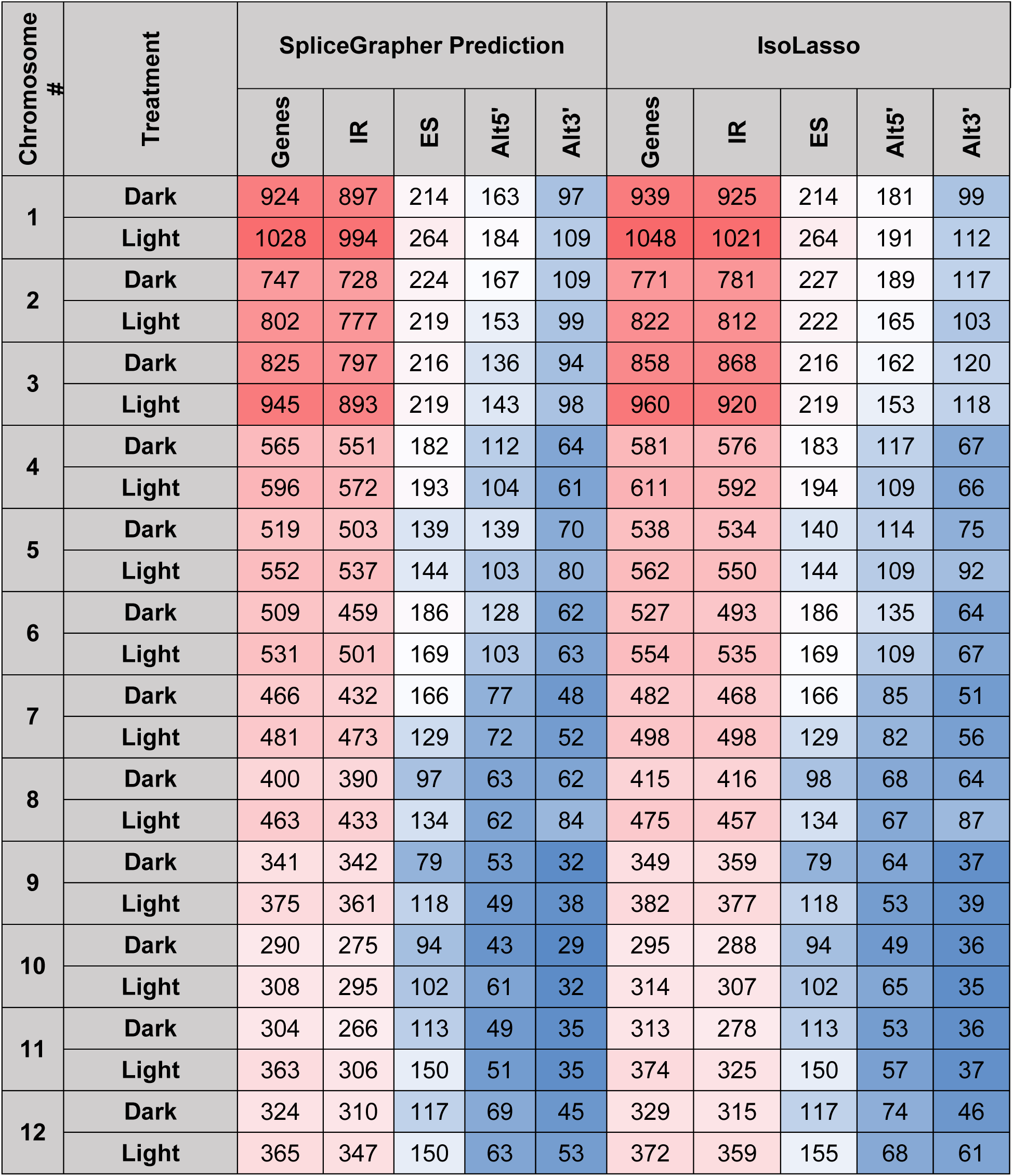
Summary of spliced genes and events under dark and light conditions detected by the SpliceGrapher and the IsoLasso workflows. IR: Intron retention; ES: Exon Skipping; Alt5’: alternative 5′ splice site; Alt3’: alternative 3′ splice site. Color scales: Red (Large counts) to Blue (Lower counts).

In our differentially expressed and spliced genes (DES), we found that approximately 30% (4369) differentially expressed (DE) genes also undergo splicing (AS). Approximately 10% (2440) of the non-DE genes undergo splicing (Figure 1). Light-upregulated DES genes show enrichment for various molecular functions like oxidoreductase, hydrolase, isomerase activity, and RNA binding, which play roles in photosynthesis, transmembrane transport, small molecule metabolic process, and located in the cellular components, chloroplast, and membrane-bound organelles. In contrast, the light-downregulated DES genes enriched for RNA processing, response to stress, protein binding, and macromolecule metabolic process, and localized in cellular components cytoplasm and nucleus (Supplementary Figure S4).

We also surveyed the DES aspects of the 57 known spliced genes in rice ^25,62^ and observed splicing in 41 genes (Table 3). Of these, seven spliced under dark conditions, three under light conditions, and 31 genes spliced under both dark and light conditions. Of the known genes, we also observed that 18 genes were light-upregulated and 24 downregulated. Only 32 genes showed a DES profile. Three light-upregulated genes (Os02g0130600, Os05g0348100, Os12g0567300) show complete splicing in light compared to intron retention in the dark. In contrast, the two light-downregulated genes (Os02g0666200 and Os04g0656100) show intron retention.

**Table 3:** Expression and splicing profile of genes in our data that are known to undergo splicing from previous reports.

| Known spliced genes | Spliced in | Regulation in light | Gene description |
| --- | --- | --- | --- |
| Os01g0619000 | Dark, Light | Down | RNA recognition motif domain-containing protein |
| Os01g0649900 | Dark, Light | - | GDSL esterase/lipase protein 20 |
| Os01g0764000 | Dark, Light | Up | PHI glutathione s-transferase 2 |
| Os01g0834400 | Dark | - | HAP3A subunit of CCAAT-box binding complex |
| Os02g0122800 | Dark, Light | Down | Similar to Arginine/serine-rich splicing factor |
| Os02g0130600 | Dark | Up | Conserved hypothetical protein |
| Os02g0161900 | Dark, Light | Down | Rice ubiquitin2 |
| Os02g0197900 | Dark, Light | Down | OsSTA53 |
| Os02g0274900 | Dark, Light | Up | Similar to metabolite transport protein CSBC |
| Os02g0577100 | Dark, Light | Down | Zinc finger, RING/PHD-type domain containing protein |
| Os02g0666200 | Light | Down | Aquaporin |
| Os02g0834000 | Dark, Light | Up | Rac-like GTP-binding protein 5 |
| Os02g0291000 | - | Down | Calcineurin B-like protein 7 |
| Os02g0291400 | - | - | Calcineurin B-like protein 8 |
| Os02g0823100 | - | Up | Plasma membrane intrinsic protein 1;3 |
| Os03g0265600 | Dark, Light | Down | Transformer-2-like protein |
| Os03g0314100 | Dark, Light | Down | DEAD-like helicase |
| Os03g0395900 | Dark, Light | Down | Splicing factor |
| Os03g0670700 | Dark, Light | Down | Glycine-rich rna-binding protein 3 |
| Os03g0698500 | Dark, Light | Down | Similar to Yippee-like protein 3 |
| Os03g0745000 | Dark, Light | - | Heat stress transcription factor A-2a |
| Os03g0717600 | - | Down | Zinc finger, C2H2-type |
| Os04g0115400 | Dark, Light | Down | D111/G-patch domain containing protein |
| Os04g0649100 | Dark, Light | - | Shattering abortion 1 |
| Os04g0656100 | Light | Down | H<sup>+</sup>-ATPase |
| Os04g0665800 | Dark, Light | Up | Similar to H1005F08.12 protein |
| Os04g0402300 | - | - | Cysteine-type peptidase |
| Os04g0479200 | - | Up | Similar to NAD-dependent isocitrate dehydrogenase c;1 |
| Os05g0348100 | Dark | Up | Similar to CRR23 (chlororespiratory reduction 23) |
| Os05g0463800 | Dark, Light | Down | HAP3C subunit of CCAAT-box binding complex |
| Os05g0554400 | Dark, Light | Down | Phosphatidyl serine synthase family protein |
| Os05g0574700 | Dark, Light | - | Similar to cDNA clone:002-182-C01 |
| Os05g0534400 | - | Up | Calcineurin B-like protein 4 |
| Os05g0548900 | - | Up | Phosphoethanolamine N-Methyltransferase 2 |
| Os06g0172800 | Light | - | Similar to alkaline alpha galactosidase 2 |
| Os06g0651600 | Dark, Light | Down | Protein phosphatase 2C58 |
| Os06g0727200 | Dark | Down | Catalase B |
| Os06g0128500 | - | Up | Ribosomal protein L47, mitochondrial family protein |
| Os06g0133000 | - | - | Glutinous Endosperm, waxy |
| Os06g0506600 | - | - | Ubiquitin-conjugating enzyme 17 |
| Os07g0490400 | Dark, Light | Up | FK506 binding protein 20-2 |
| Os07g0574800 | - | Up | Tubulin alpha-1 chain |
| Os07g0613300 | - | - | Similar to PAUSED |
| Os08g0191600 | Dark, Light | Up | Autophagy associated gene 8C |
| Os08g0530400 | Dark, Light | Up | Moco containing protein, Similar to sulfite oxidase |
| Os08g0436200 | - | - | Zinc finger, RING/PHD-type domain containing protein |
| Os10g0115600 | Dark, Light | Down | U1 snRNP 70K |
| Os10g0535800 | Dark, Light | - | Cys-rich domain containing protein |
| Os10g0564900 | Dark | Down | Similar to protein kinase CK2 regulatory subunit CK2B2 |
| Os10g0567400 | Dark, Light | Up | Chlorophyll a oxygenase 1 |
| Os10g0577900 | Dark, Light | - | Glycerol-3-phosphate acyltransferase |
| Os10g0411500 | - | Down | Q calmodulin-binding region domain containing protein |
| Os11g0157100 | Dark, Light | Down | Cyclin-T1-4 |
| Os11g0700500 | Dark | - | MybAS1 |
| Os11g0600700 | - | Up | Zinc finger, RING domain protein |
| Os12g0567300 | Dark | Up | MybAS2, R2R3-Myb |
| Os12g0632000 | Dark, Light | Down | Glycine-rich Protein GRP162 |

### Light-mediated shift in gene expression and splicing events

#### Circadian clock pathway genes

The circadian clock is entrained by light, and the diurnal photoperiod regulated clock genes ^63,64^. We investigated the light-regulated DES profile of clock genes. Most of the circadian clock genes spliced under either or both conditions. Four genes, *Casein kinase alpha subunit* (*CK2α-1/Hd6*, Os03g0762000), *Casein kinase beta subunit* (*CK2β-2*, Os10g0564900), *Phytochrome B* (*PhyB,* Os03g0309200;), and *Timing of CAB Expression 1* (*TOC1*, Os02g0618200), showed differential splicing (only reference splicing events or complete splicing and no novel isoforms) upon light treatment compared to the presence of unspliced intron in the absence of light (Supplementary Figure S5). In contrast, only one gene, *Phytochrome Interacting Factor 3* (*PIF3*, Os01g0286100), showed intron retention under light. All five genes were upregulated in light (Supplementary Figure S5). Seven genes, *Casein kinase alpha subunit* (*CK2α-2*, Os03g0763000), *Casein kinase beta subunit* (*CK2β-1*, Os07g0495100), blue light receptor *Cryptochrome 2* (*CRY2*, Os02g0625000), *Gigantea* (*GI*, Os01g0182600), *Pseudo-Response Regulator 37* or *Heading date 2* (*PRR37*, *Hd2*, Os07g0695100), and *Pseudo-Response Regulator 73* (*PRR73*, Os03g0284100), showed differential splicing under both conditions (Supplementary Figure S5, S6). Exon skipping and alternative 5’ (Alt.5’) splicing events in the *CK2α-2* (Os03g0763000) gene were induced in light. Both *CK2β-1* (Os07g0495100) and *GI* (Os01g0182600) display incomplete splicing of intron (IR) in the dark. *PRR37* showed three types of splicing events (Alt.3’, ES, and IR) under light, whereas the second IR event observed toward the 3’ end under dark was spliced in light. Conversely, *PRR73* showed three types of events (Alt.3’, ES, and IR) under dark conditions, whereas under light conditions, two new Alt.5’ events were identified. Expression of both PRR genes (*PRR37* and *PRR73*) was light downregulated (Supplementary Figure S5). Light-upregulated *PRR95* shows an additional unspliced intron in light.

#### MVA and MEP pathway genes

Light regulates the biosynthesis and accumulation of secondary plant products like isoprenoid-derived metabolites required for plant growth and development. Isoprenoid precursors are synthesized via cytosolic MVA and plastid-localized MEP pathways, and their fluxes are regulated by light ^65,66^, and their products are essential for the successful development and function of chloroplast. We observed that MEP pathway genes were light-upregulated compared to the downregulated MVA pathway genes (Figure 2). Light positively regulates MEP pathway activity, whereas MVA pathway activity appears negatively regulated by light ^67^.

**Figure 2:**
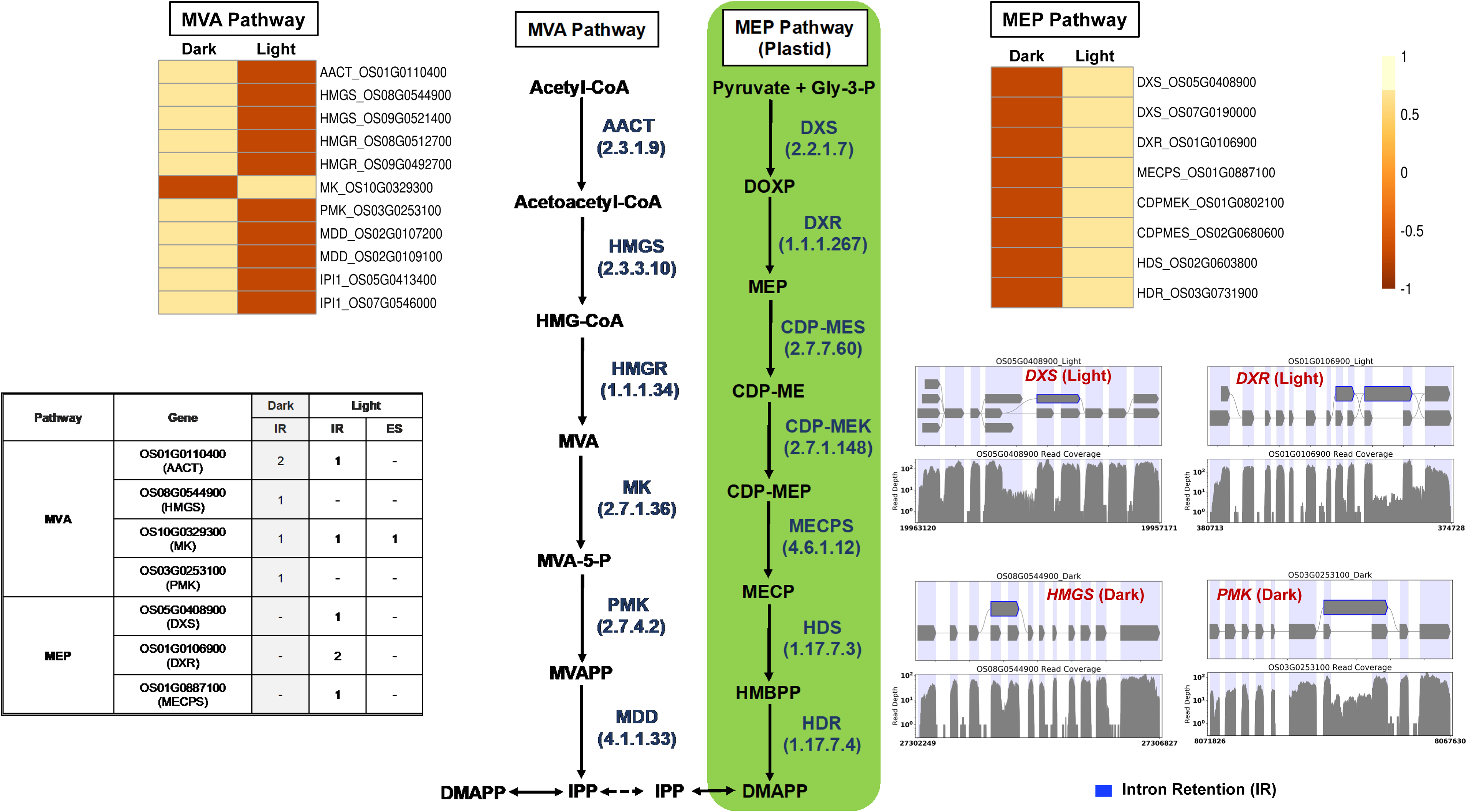
Gene expression and splicing of cytosolic mevalonate (MVA) and plastid-localized methyl erythritol 4-phosphate (MEP) pathway genes. *AACT*, acetyl-CoA acetyltransferase; *HMGS*, hydroxymethyl glutaryl CoA synthase; *HMGR*, 3-hydroxy-3-methylglutaryl-coenzyme A reductase; *MK*, mevalonate kinase; *PMK*, phosphomevalonate kinase; *MDD*, mevalonate diphosphosphate decarboxylase; *DXS*, 1-deoxy-D-xylulose-5-phosphate synthase; *DXR*, 1-deoxy-D-xylulose-5-phosphate reductoisomerase; *CDPMES*, 2-C-methyl-D-erythritol4-phosphate cytidylyl transferase; *CDPMEK*, 4-diphosphocytidyl-2-C-methyl-D-erythritol kinase; *MECPS*, 2-C-methyl-D-erythritol2,4-cyclodiphosphate synthase; *HDS*, 4-hydroxy-3-methylbut-2-enyldiphosphate synthase; *HDR*, 4-hydroxy-3-methylbut-2-enyldiphosphate reductase. Blue: intron retention, Green: exon skipping, Orange: Alt3’ splicing, Purple: Alt5’ splicing.

We investigated the light-induced differential splicing events in the MEP and MVA pathway genes. Four MVA pathway genes (*acetyl-CoA acetyltransferase: AACT, hydroxymethyl glutaryl CoA synthase: HMGS, mevalonate kinase: MK, phosphomevalonate kinase: PMK*) and 3 MEP pathway genes (*1-deoxy-D-xylulose-5-phosphate synthase: DXS, 1-deoxy-D-xylulose-5-phosphate reductoisomerase: DXR, 2-C-methyl-D-erythritol2,4-cyclodiphosphate synthase: MECPS*) showed intron retention events in dark (Figure 2). Gene *MK* showed a novel light-induced exon-skipping event.

### lncRNA discovery and splicing during photomorphogenesis

lncRNAs are known to regulate diverse biological processes in plants, including photomorphogenesis ^68^, and as described above, are categorized into four major types based on their location in the gene space and on the DNA strand. We used our transcriptome data to characterize light-induced rice lncRNAs (Supplementary Figure S7). We identified 1485 and 1407 lncRNA transcripts in the dark and light samples, of which 309 were common. Many lncRNAs from intergenic and intronic regions were identified (Supplementary Figure S7). To discover novel lncRNAs, the transcripts from our datasets were queried against the custom BLAST database of rice lncRNAs. About 44% of the total lncRNA transcripts (2583) found a match in the BLAST database (Supplementary Table S6). 173 lncRNAs spliced under dark and 142 spliced under light conditions, most located in the intergenic regions (Supplementary Figure S7). Interestingly, all the spliced lncRNA genes from both datasets were from Chr1, Chr10, Chr11, and Chr12. The maximum number of spliced lncRNA genes was from Chr1, followed by Chr11, 12, and 10. Among all the splicing events, IR was predominant under both conditions.

We found lncRNA transcripts from light (TCONS_00012152) and dark (TCONS_00011794) datasets that were present on the antisense strand overlapping an exon region of the *MADS27* (Os02g0579600) gene. The transcript from the light-treated sample appears longer and showed two alternative 3’ splice sites and one exon skipping event. The TCONS_00011794 may compete with MADS27 to form a complex with miRNA osa-miR444a known to play multiple roles in the nitrate-dependent development pathway ^69^. Among the list of lncRNAs present on the antisense strand overlapping annotated genes, we identified a light-induced lncRNA (TCONS_00001515) overlapping *ERF99* transcription factor (Os01g0868000) exon that may play a role in its silencing. *ERF99* plays a central role in mediating abiotic stress responses ^70^. Similarly, lncRNA transcript (TCONS_00015825) is present exclusively in light is transcribed from the intronic region of the antisense strand of a B-type response regulator gene (Os03g0224200), known for its involvement in cytokinin signaling, meristem maintenance, and stress response ^71,72^. The lncRNA transcript (TCONS_00026617) present exclusively in the dark overlaps the intronic region of a phytochrome-interacting factor gene *PIF14* (Os07g0143200). *PIF14* is known to bind the active form of phytochrome B and plays a crucial role in cross-talk between light and stress signaling ^73^.

## Discussion

Light is known to induce changes in the transcriptome, metabolome, and proteome of the plants ^74^, which not only regulates the development, function, and physiology of the chloroplast but also provides signals for modulating the plant’s morphological, developmental, and physiological adaptions in response to growth environment ^8,40,75–77^. Though plants experience constantly changing light conditions under the natural environment daily during their lifetime, early development from germination to the seedling stage and acquiring the full autotrophic capability is very much programmed by exposure to light. Therefore, we investigated transcriptome modulation in rice seedlings during photomorphogenesis. Compared to earlier studies in rice and *Arabidopsis*, where ~20% of the genes were reported differentially expressed in dark-grown etiolated seedlings compared to light-exposed green seedlings ^9,78,79^, we observed that the transition from skotomorphogenesis to photomorphogenesis alters differential expression of ~38% of the rice genes.

Functional annotation of the differentially expressed genes enriched for roles in secondary metabolism, chloroplast-related biosynthetic pathways, hormone biosynthesis and signaling, amino acid biosynthesis, fatty acid metabolism, *etc*. (Supplementary Figure S3, Supplementary Table S3). This result was expected based on the earlier reports on light regulation of such events ^24,80^.

The plastids develop into chloroplasts by following an essential step of developing the thylakoid membrane system and recruiting and assembling components of light and dark reactions to establish a functional photosynthetic process. Many terpenoid compounds are essential to plants’ light-harvesting function and protect against damage from reactive oxygen species (ROS). The basic isoprenoid units for terpenoid biosynthesis, such as tocopherols, plastoquinones, carotenoids, chlorophylls, and precursors of the growth hormones gibberellins and abscisic acid, are synthesized by the plastid-localized MEP pathway ^81,82^. At the same time, isoprenoid units synthesized by the MVA pathway contribute to the synthesis of triterpenes, phytosterols, and phytohormones ^83^. MEP and MVA pathways complement and contribute to the biosynthesis of chlorophylls and carotenoids required for plastid development ^84^. MEP pathway genes were light-upregulated in our transcriptome data compared to the light-downregulated MVA pathway genes (Figure 2). These results were consistent with earlier reports ^85,86^ and confirm that light acts as a critical regulator in modulating the availability of isoprenoid precursors during photomorphogenesis ^87^. Arabidopsis circadian clock genes (*LHY*, *PRR9*, *CCA1*, *TOC1*) regulate MVA and MEP pathways ^87^. *AtTOC1* is also known to regulate the G1-to-S phase transition in the mitotic cell cycle in early leaf development ^88^. In our dataset, we did not observe any significant change in the expression of *LHY/CCA1*(Os08g0157600), *PRR95* (Os09g0532400, a homolog of *AtPRR9*), and *TOC1* (Os02g0618200). However, all three genes spliced differently. Under the light, *TOC1* showed no splicing but retained two introns in the dark. We observed one exon-skipping event in light of the *LHY/CCA1* gene, whereas *PRR95* showed one additional intron retention (Supplementary Figure S5). Similar to *Arabidopsis* ^88^, the functional enrichment analysis also identified the *TOC1*-regulated DNA biosynthesis pathway of the G1-to-S transition of mitotic cell cycle (Figure 3) along with a greater coverage of reaction events in the light-upregulated set. To our knowledge this is the first report of the light regulation of mitotic cell-cycle pathway in rice.

**Figure 3:**
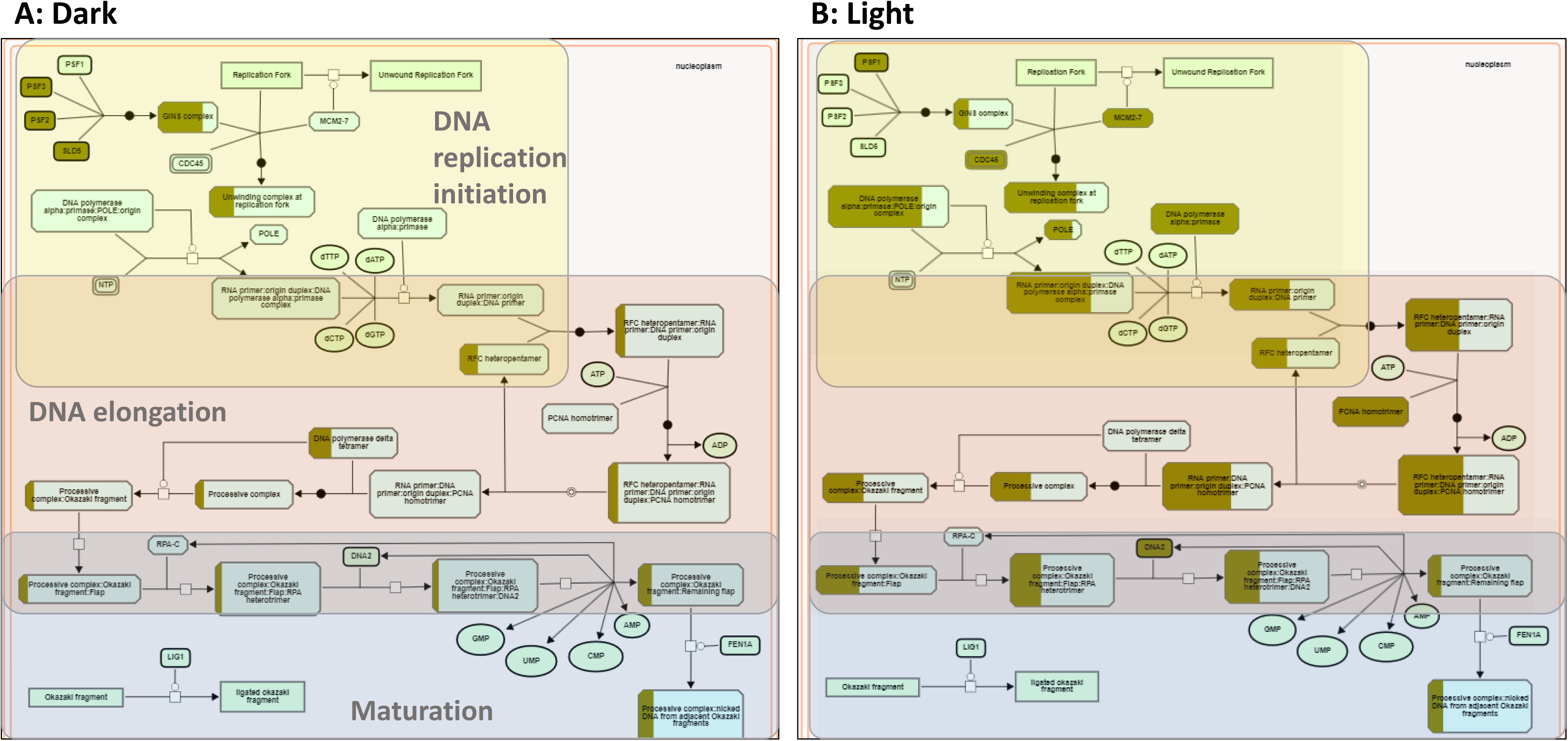
Plant Reactome pathway enrichment showing light-regulated genes and reaction event coverage for the DNA biosynthesis pathway of the G1-to-S phase mitotic cell cycle.

Previous transcriptome studies showed that ~42% of *Arabidopsis*, ~63% of *soybean*, and ~56% of maize genes undergo alternative splicing events ^89–91^. Studies on rice by Wang and Brendel ^25^ reported IR for 69% of 6568 AS genes, and Campbell *et al.* ^92^ reported 44.7% of 8772 AS genes. In our dataset, 4369 genes undergo differential expression and alternative splicing, called DES genes (Figure 1A). About 80% (5395) of the spliced genes (6809) showed IR events, suggesting that IR is the major splicing event. The more significant number of splicing events in the light-treated samples suggests that light-mediated gene expression and post-transcriptional mRNA splicing play an important role in photomorphogenesis. The splicing events in rice circadian clock genes suggest that IR regulates the expression of PRR genes (Supplementary Figure S5). Many of the light-regulated genes of the MEP and MVA pathways show IR events, suggesting that light plays a positive role in completing the splicing and provides a hypothesis that exposure to light is an adaptive feature and even if the gene is expressed in dark, it stays unspliced until it encounters light. Our data showed the presence of all three states, partially spliced in the dark (*GI*, Os01g0182600), fully spliced in light (*GI*, Os01g0182600), and both states in light (*CK2β-1*, Os07g0495100) (Supplementary Figure S6).

lncRNAs play regulatory functions in essential biological processes such as vernalization, photomorphogenesis, and stress regulation ^39,93,94^ and display splicing ^95^. 6480 long intergenic non-coding RNAs (lincRNAs) were identified from 200 *Arabidopsis* transcriptome datasets ^96^; however, we identified 827 lincRNAs in the dark and 727 lincRNAs in the light datasets. Corona-Gomez and coworkers ^97^ characterized the splicing conservation of lncRNAs in *Brassicaceae*, which revealed that ~18% of lincRNAs display splicing conservation in at least one exon. In our dataset, ~10% of lncRNA transcripts undergo splicing, and the majority were of long intergenic non-coding RNA type (Supplementary Figure S7). Komiya and colleagues reported ^98^ that phased small interfering RNAs (phasiRNAs) are generated from over 700 lincRNAs, and these phasiRNAs bind to rice argonaute protein MEL1. MEL1 has a specific function in developing pre-meiotic germ cells and the progression of meiosis. In our dataset, many lincRNAs transcribed from MEL1-phasiRNA clusters appeared.

Identification of light-induced lncRNAs antisense to *ERF99*, B-type response regulator genic region may play a role in transcriptional regulation. *ERF99* is known to modulate root architecture and downregulated in crown root primordia ^99^. *MADS27* gene targeted by lncRNAs present in both dark and light conditions. However, the dark lncRNA (TCONS_00011794) is longer and, based on its predicted RNA fold structure and lower free energy, is likely more stable than its transcript TCONS_00012152 in light. For the MADS27 gene, we also observed an additional IR event in light. MADS27-miR444 complex is known to play a role in plant development in a nitrate-dependent manner ^100–102^. Therefore, we investigated the nitrogen assimilation pathway genes. Nitrogen assimilation is necessary for sustaining plant growth and development. Various nitrogen assimilatory enzymes are known to show isoform and cellular component-specific responses under light and dark conditions ^103,104^. We observed light upregulation for the nitrate transporter, nitrate reductase, and nitrite reductase genes. We also observed differential splicing patterns for these genes (Supplementary Figure S8). We also found light regulation of the recently reported rice abiotic and biotic stress response genes ^105,106^.

## Conclusion

This study suggests that light is a significant regulatory factor controlling genome-wide gene expression through an alternative splicing mechanism in rice. All spliced genes did not necessarily produce novel isoforms, which indicates the coupling of AS and nonsense-mediated decay (NMD). NMD prevents the translation of mutant mRNAs harboring potential premature stop codons by targeting them for degradation. In *Arabidopsis*, 77.2% of light-regulated AS events exhibit NMD features within a splicing isoform ^107^. We conclude that light induces a significant number of splicing events in rice protein-coding and non-coding lncRNA genes. This photomorphogenesis transcriptome study is a valuable resource for lncRNA research in rice and provides insights into the portion of the genome regulated at the level of alternative splicing in response to light. We expect the condition-dependent novel splice events discovered in this study will help improve the annotation of the rice reference genome and pan-genomes ^49,108^

## Availability of data

The sequence reads and metadata were deposited at EMBL-EBI ArrayExpress (accession number E-MTAB-5689).

## Supplementary material

**Supplementary Figure S1:**

Expression pattern of differentially expressed genes (14,766) with target FDR controlled at 5%

**Supplementary Figure S2:**

Gene Ontology enrichment analysis of differentially expressed genes.

**Supplementary Figure S3:**

Pathway enrichment analysis using the Plant Reactome. **(A)** Plant Reactome pathway enrichment analysis plots, **(B)** Unique and shared pathways enriched for the light upregulated and downregulated gene sets; **(C)** Counts of genes mapped to some of the common Plant Reactome pathways. Light upregulated (green) and downregulated (grey).

**Supplementary Figure S4:**

Bar plots of most significantly enriched GO terms for DES genes with −log10 transformed FDR values. Light-upregulated (green), light-downregulated (grey).

**Supplementary Figure S5:**

The summary of transcript splicing events and expression pattern of circadian clock genes during morphogenesis.

**Supplementary Figure S6:**

Transcript graphs showing splicing of *Casein kinase beta subunit:CK2β-1* (Os07g0495100), *Gigantea:GI* (Os01g0182600) genes during photomorphogenesis.

**Supplementary Figure S7:**

Classification of lncRNAs identified in dark and light.

**Supplementary Figure S8:**

An overviw of splicing and expression pattern of nitrogen assimilation cycle genes during photomorphogenesis. NRT: Nitrate transporter; NR: Nitrate reductase; NiR: Nitrate reductase.

**Supplementary Table S1:**

Summary of the RNA-Seq read mapping to the reference rice genome.

**Supplementary Table S2:**

Differentially expressed genes

**Supplementary Table S3:**

Plant Reactome Pathway enrichment analysis for light-regulated genes

**Supplementary Table S4:**

Transcription factor genes regulated in light

**Supplementary Table S5:**

Transcription factors regulating MVA pathway genes

**Supplementary Table S6:**

lncRNA annotation using BLAST

