## Supplementary material for "Transcriptional modulation during photomorphogenesis in rice seedlings": Supplementary Figure S1.pdf

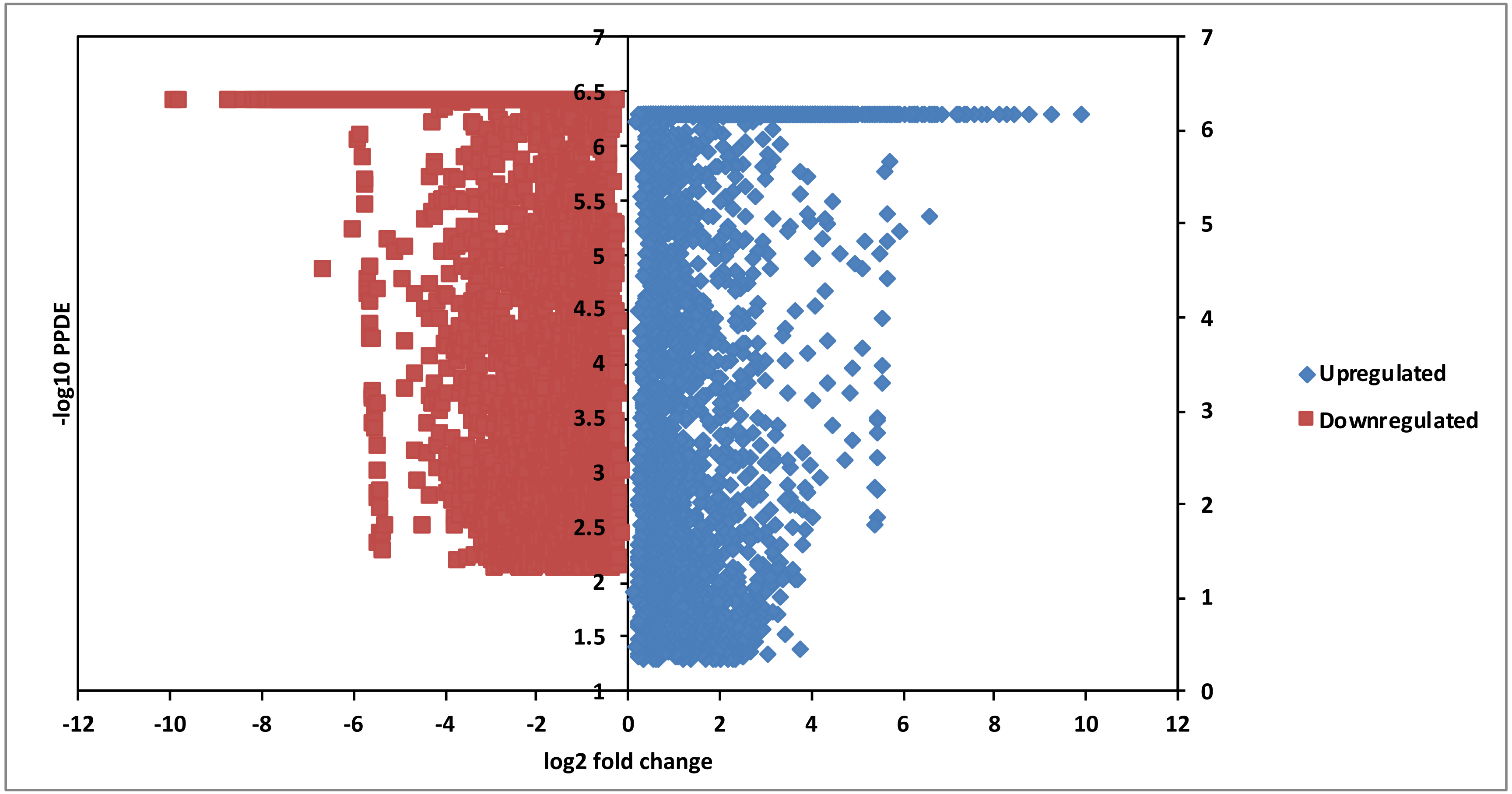


**Supplementary Figure S1:** Expression pattern of differentially expressed genes (14,766) with target FDR controlled at 5%
