## Supplementary material for "Transcriptional modulation during photomorphogenesis in rice seedlings": Supplementary Figure S2.pdf

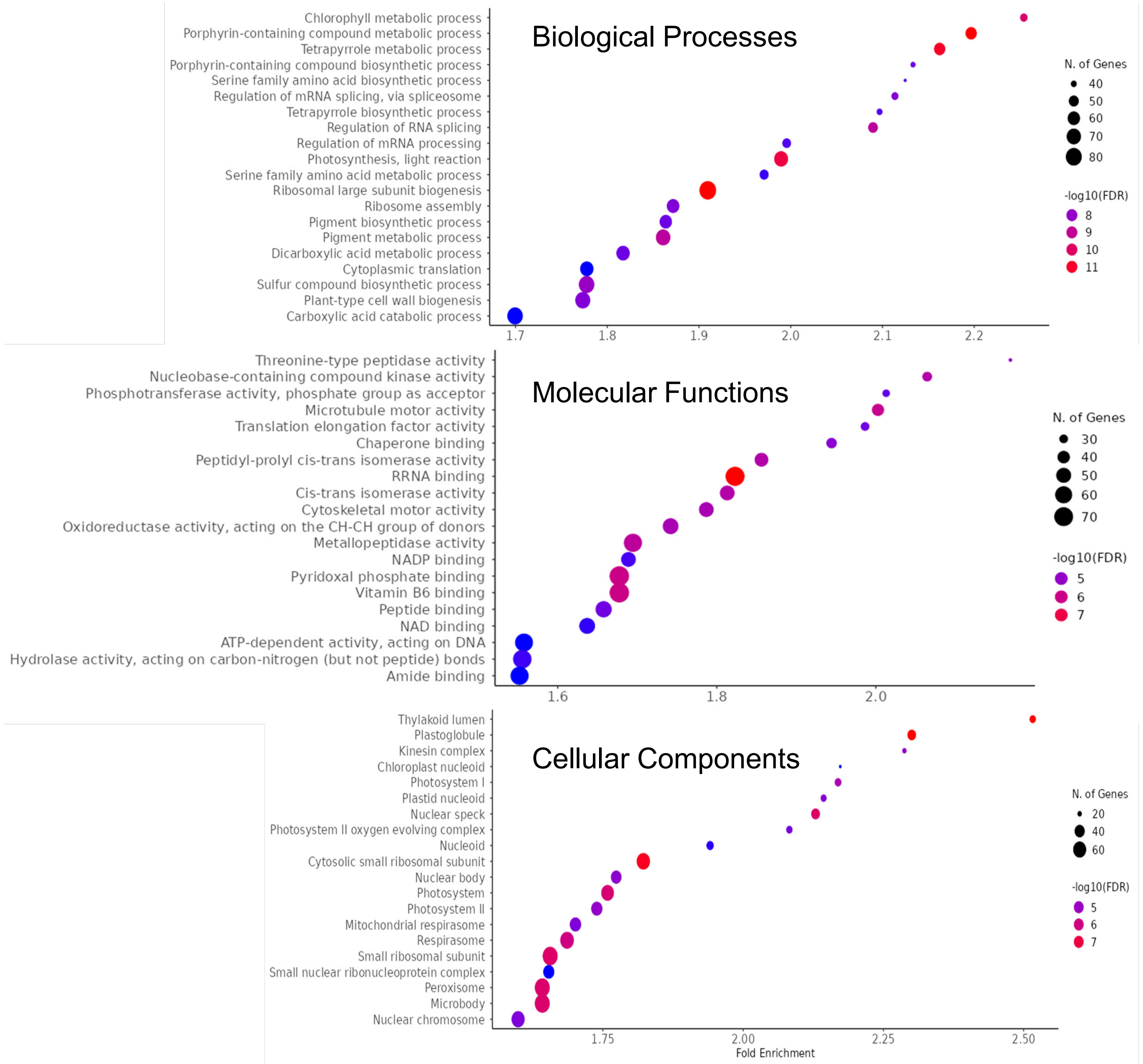


**Supplementary Figure S2:** Gene Ontology enrichment analysis of the differentially expressed genes.
