## Supplementary material for "Transcriptional modulation during photomorphogenesis in rice seedlings": Supplementary Figure S3.pdf

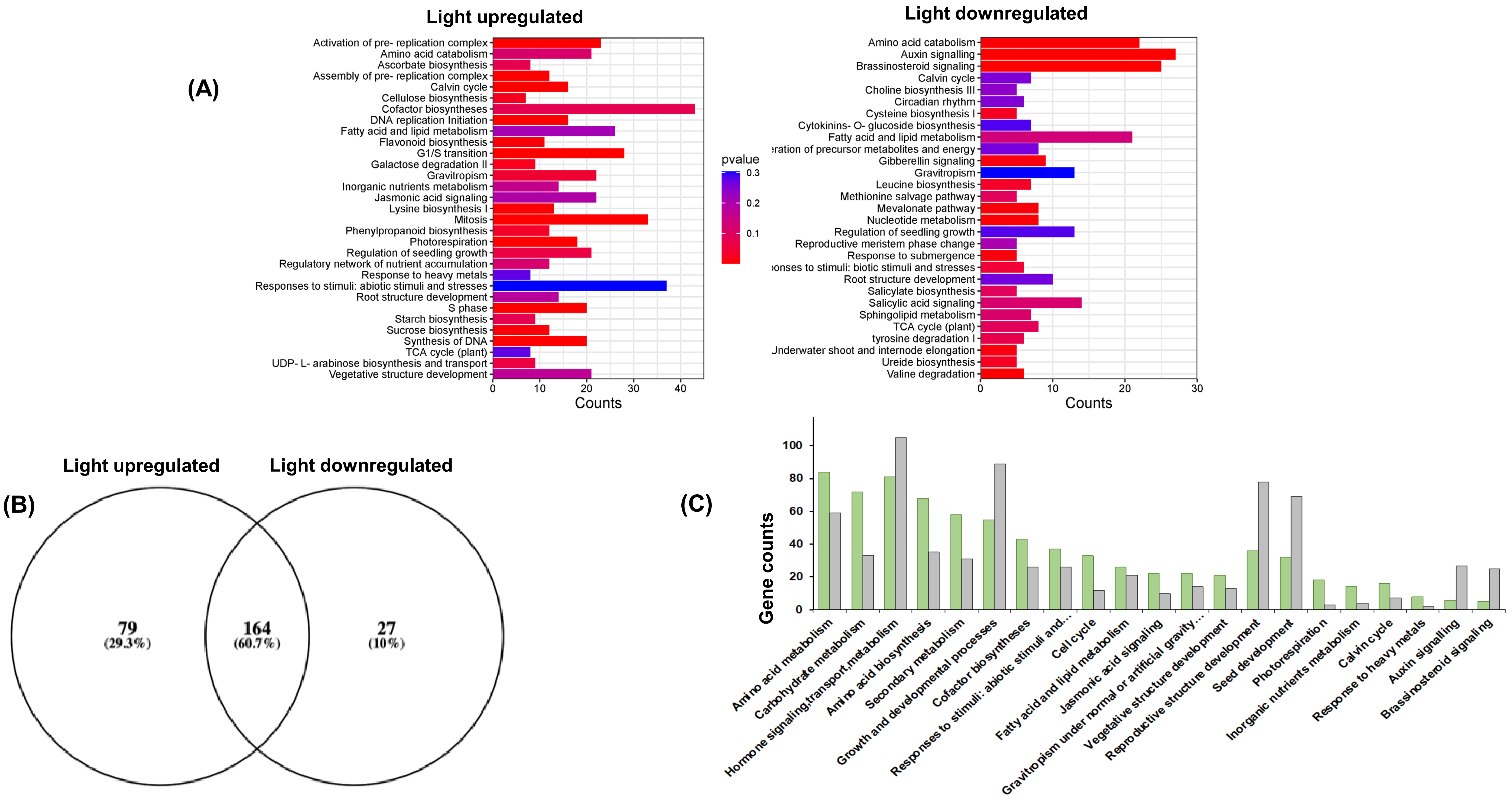


**Supplementary Figure S3:** Pathway enrichment analysis using the Plant Reactome. (A) Plant Reactome pathway enrichment analysis plots, **(B)** Unique and shared pathways enriched for the light upregulated and downregulated gene sets; **(C)** Counts of genes mapped to some of the common Plant Reactome pathways. Light upregulated (green) and downregulated (grey).
