## Supplementary material for "Transcriptional modulation during photomorphogenesis in rice seedlings": Supplementary Figure S4.pdf

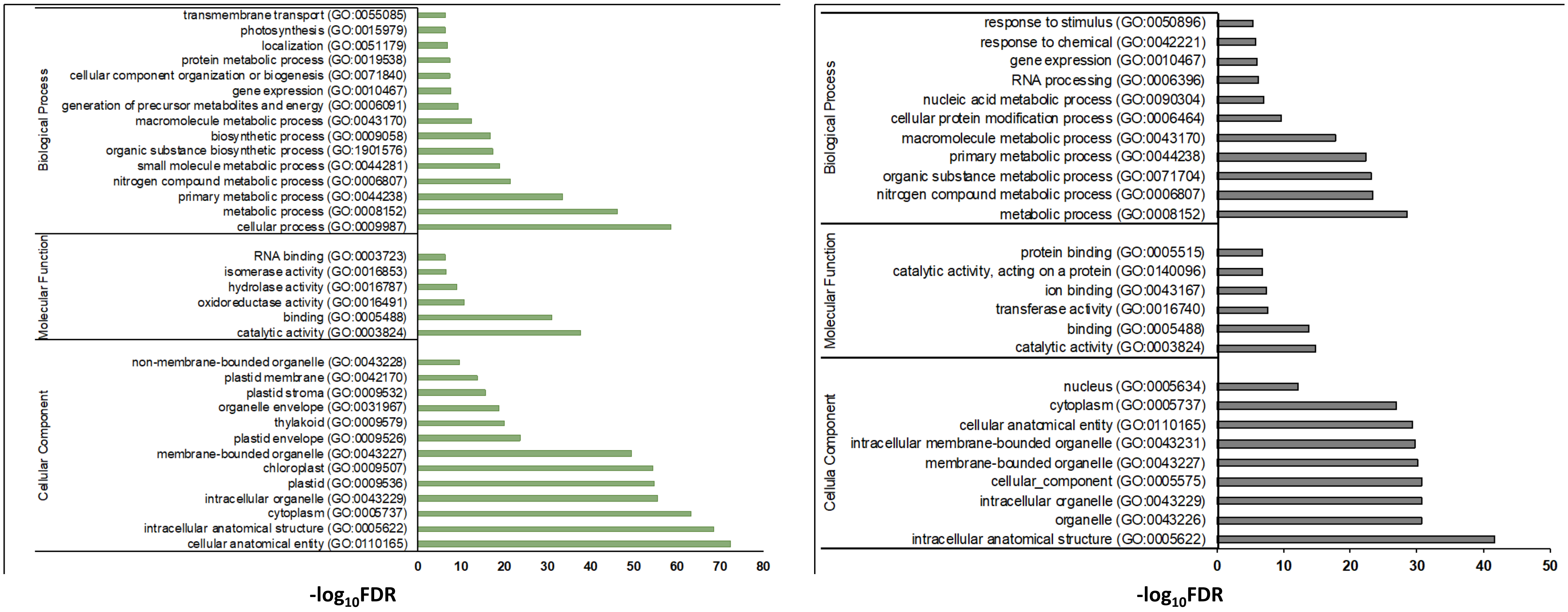


**Supplementary Figure S4**: Bar plots of most significantly enriched GO terms for DES genes with -$\log_{10}$ transformed FDR values. Light-upregulated (green), light-downregulated (grey).
