## Supplementary material for "Transcriptional modulation during photomorphogenesis in rice seedlings": Supplementary Figure S5.pdf

| Genes | Light | | | | Dark | | | |
| --- | --- | --- | --- | --- | --- | --- | --- | --- |
| | IR | ES | Alt5’ | Alt3’ | IR | ES | Alt5’ | Alt3’ |
| CK2α-1 (Os03g0762000) | - | - | - | - | 1 | - | - | - |
| CK2α-2 (Os03g0763000) | 1 | 1 | 1 | - | 1 | - | - | - |
| CK2β-2 (Os10g0564900) | - | - | - | - | 1 | - | - | - |
| PhyB (Os03g0309200) | - | - | - | - | 1 | - | - | - |
| TOC1 (Os02g0618200) | - | - | - | - | 2 | - | - | - |
| PIF3 (Os01g0286100) | 1 | - | - | - | - | - | - | - |
| CRY2 (Os02g0625000) | 2 | - | - | - | 1 | - | - | 1 |
| PRR37 (Os07g0695100) | 3 | 1 | - | 1 | 2 | - | - | - |
| PRR73 (Os03g0284100) | 3 | - | 2 | - | 2 | 1 | - | 3 |
| PRR95 (Os09g0532400) | 2 | - | 1 | - | 1 | - | 1 | - |

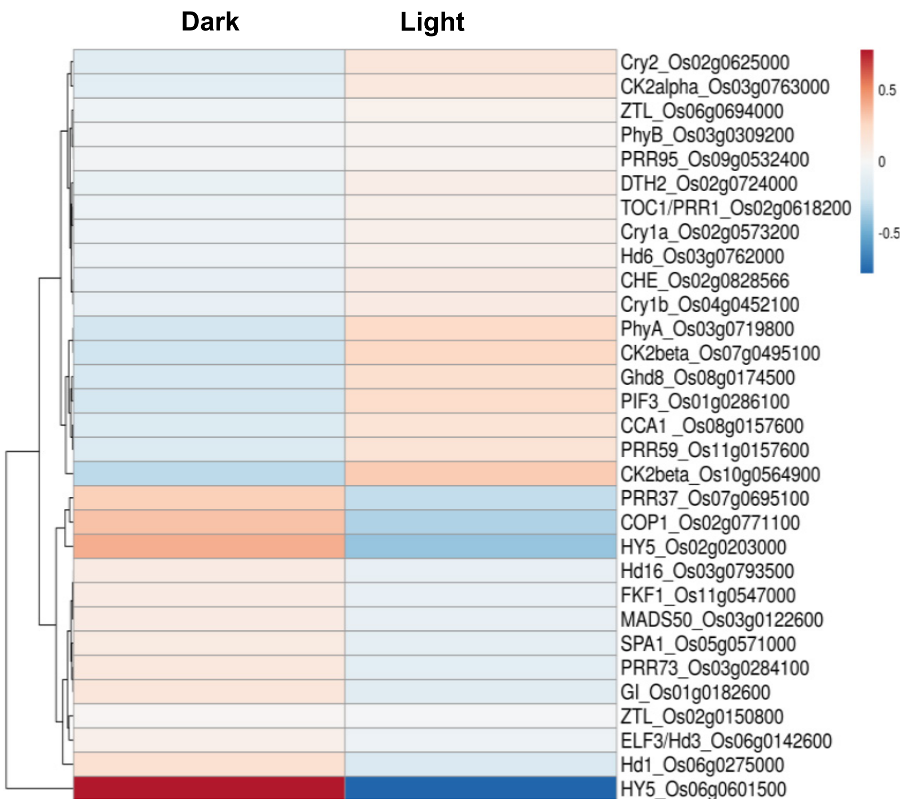


**Supplementary Figure S5:** Splicing events and expression pattern of circadian clock genes under dark and light conditions.
