## Supplementary material for "Transcriptional modulation during photomorphogenesis in rice seedlings": Supplementary Figure S6.pdf

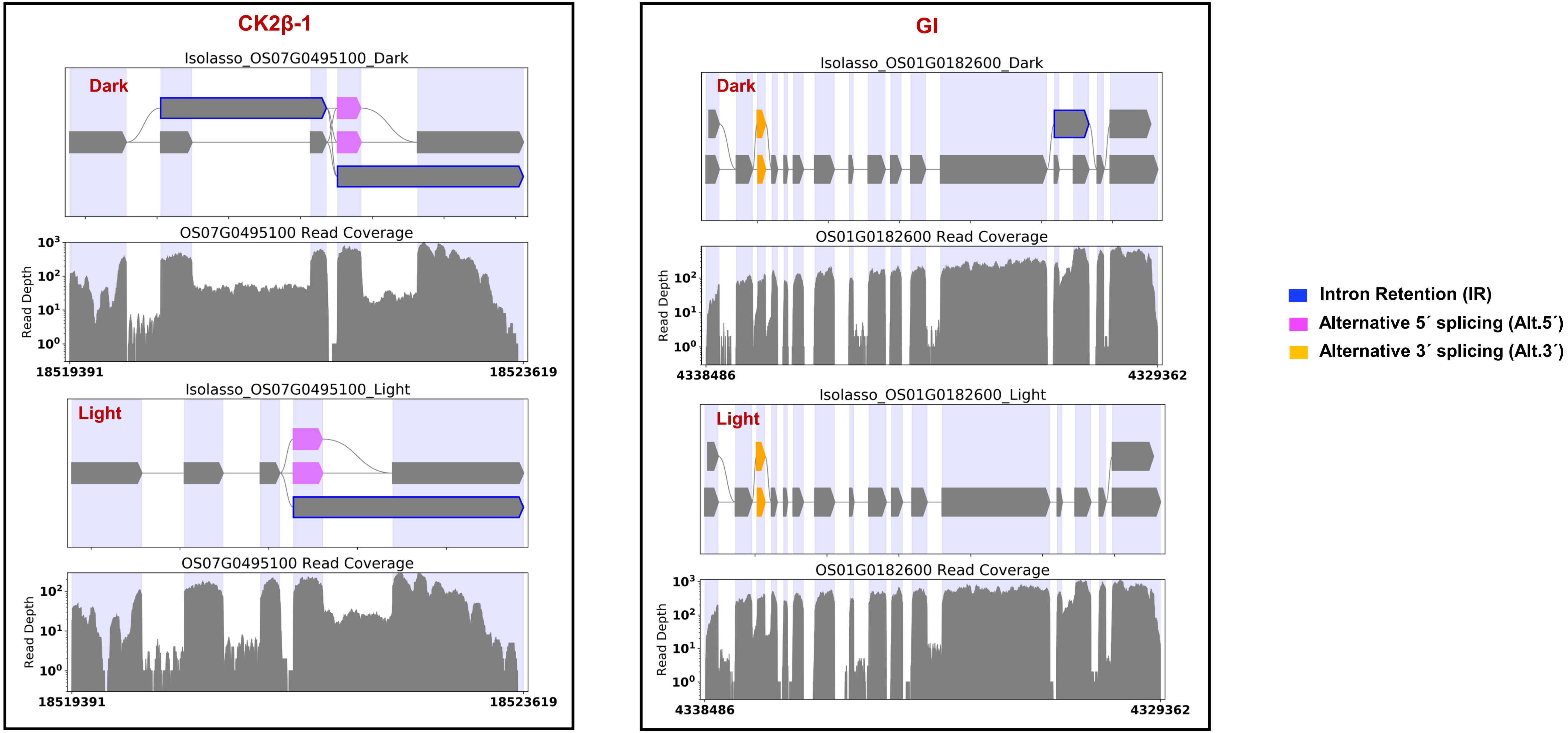


**Supplementary Figure S6:** Splicing of *Casein kinase beta subunit:CK2β-1* (Os07g0495100), *Gigantea:GI* (Os01g0182600) genes under dark and light conditions.
