## Supplementary material for "Transcriptional modulation during photomorphogenesis in rice seedlings": Supplementary Figure S8.pdf

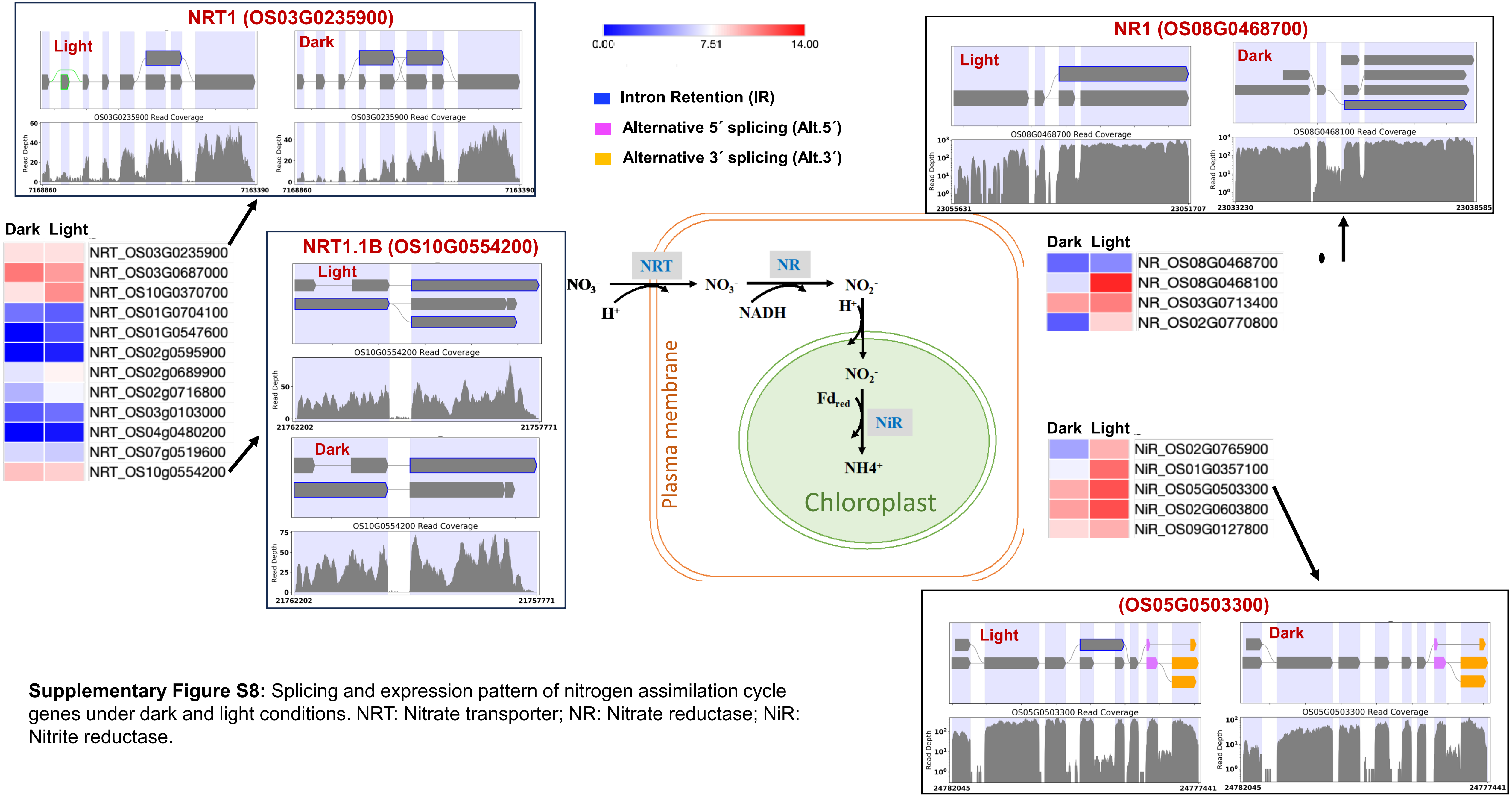


**Supplementary Figure S8:** Splicing and expression pattern of nitrogen assimilation cycle genes under dark and light conditions. NRT: Nitrate transporter; NR: Nitrate reductase; NiR: Nitrite reductase.
