## Supplementary material for "Transcriptional modulation during photomorphogenesis in rice seedlings": Supplementary Table S1.pdf

**Supplementary Table 1**: Read mapping summary of rice seedlings grown in continuous dark and exposed to light.

| Sample | Input reads | High quality reads | Reads mapped to genome |
|---|---|---|---|
| Dark-Rep1 | 13,760,553 | 13,001,563 | 12,249,629 |
| Dark-Rep2 | 14,147,179 | 13,335,860 | 12,324,418 |
| Dark-Rep3 | 13,442,449 | 12,505,262 | 11,783,274 |
| Light-Rep1 | 14,554,961 | 13,625,166 | 12,677,035 |
| Light-Rep2 | 14,577,314 | 13,755,841 | 13,040,444 |
| Light-Rep3 | 13,333,010 | 12,695,151 | 12,058,127 |
