## Supplementary figures and images for "Transcriptional modulation during photomorphogenesis in rice seedlings"

### Supplementary Figure S7.pdf

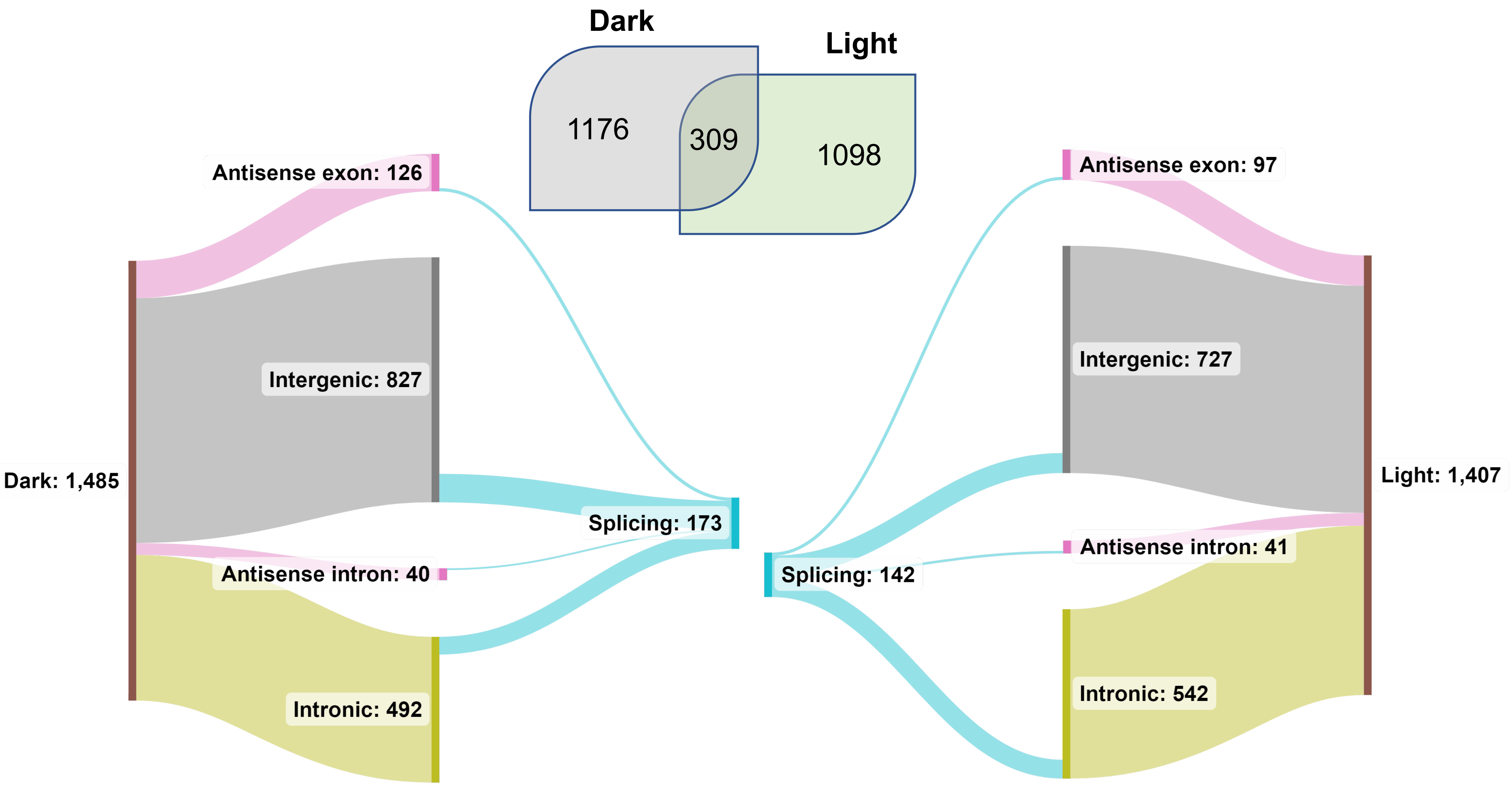


**Supplementary Figure S7:** Classification of lncRNAs identified in dark and light.
